# ROS-based lethality of *C. elegans* mitochondrial electron transport mutants grown on *E. coli* siderophore iron release mutants

**DOI:** 10.1101/707968

**Authors:** J. Amaranath Govindan, Elamparithi Jayamani, Gary Ruvkun

**Affiliations:** Department of Molecular Biology, Massachusetts General Hospital, Boston, MA 02114, USA; Department of Genetics, Harvard Medical School, Boston, MA 02115, USA

**Keywords:** *C. elegans*, *E. coli*, micronutrients, diet, reactive oxygen species, iron, siderophore, metabolism

## Abstract

*C. elegans* consumes bacteria which can supply essential vitamins and cofactors especially for mitochondrial functions ancestrally related to bacteria. Therefore, we screened the Keio *E. coli* knockout library for mutations that induce a *C. elegans* mitochondrial damage response gene. We identified 45 *E. coli* mutations that induce a the *C. elegans hsp-6::gfp* response gene. Surprisingly, four of these *E. coli* mutations that disrupt the import or removal of iron from the bacterial siderophore enterobactin were lethal in combination with *C. elegans* mutations that disrupt particular iron-sulfur proteins of the electron transport chain. Bacterial mutations that fail to synthesize enterobactin are not synthetic lethal with these *C. elegans* mitochondrial mutants; it is the enterobactin-iron complex that is lethal in combination with the *C. elegans* mitochondrial mutations. Antioxidants suppress this inviability, suggesting that reactive oxygen species (ROS) are produced by the mutant mitochondria in combination with the bacterial enterobactin-iron complex.

**Significance Statement:** The animal mitochondrion has a bacterial origin and continues to have a dialogue with the bacterial metabolisms of their microbiome. We identified 45 *E. coli* gene disruptions that induce a *C. elegans* mitochondrial damage response gene. Four of these *E. coli* mutations that disrupt the import or retrieval of iron from the siderophore enterobactin were synthetic lethal with *C. elegans* mitochondrial mutants. Antioxidants strongly suppressed the inviability of *C. elegans* mitochondrial mutants grown on the *E. coli* enterobactin siderophore utilization or import mutants. We hypothesize that reactive oxygen species are produced by C. elegans mitochondrial mutations and that this non-lethal ROS triggers ferric-chelated enterobactin to induce dramatically increased ROS, which leads to lethality.

## Introduction

*Caenorhabditis elegans*, like many nematode species, consumes bacteria which supplies many nutritional needs. *C. elegans* in the laboratory consumes *E. coli*, but in its natural habitat of rotting fruit, hundreds of bacterial species are consumed by *Caenorhabditis elegans* (1, 2). These diverse bacteria supply micronutrients to the animal such as vitamins and cofactors. The bacterial supply of such cofactors is so dependable that *C. elegans* is a heme auxotroph fully dependent on the bacteria it consumes to acquire this cofactor for many mitochondrial proteins in the electron transport chain (3). Because many of the eukaryotic nuclearly encoded proteins that localize to the mitochondria share a common ancestor with bacteria, we reasoned that other bacterial gene pathways that animals may depend upon may function for mitochondrial biogenesis or function. For example, *E. coli* mutations in the electron transport chain cytochrome *bo* terminal oxidase A gene *cyoA* cause induction of a *C. elegans* mitochondrial unfolded protein response gene, *hsp-6*, in wild type *C. elegans* (4). Other bacterial biosynthetic pathways on which the mitochondrion may still depend could be discovered using an animal reporter gene for mitochondrial dysfunction and a comprehensive bacterial collection of mutations. Therefore, we conducted a genome wide screen for *E. coli* mutations that induce the UPR^mt^ or affect the viability or growth of *C. elegans*. In this screen, we identified mutations in 45 *E. coli* genes out of more than 4000 *E. coli* gene disruptions tested that strongly activate the mitochondrial unfolded protein response gene, *hsp-6*; 24 of these 45 mutations also slow *C. elegans* growth. Because each of the 45 *E. coli* mutations induce a mitochondrial stress response, we tested for genetic interactions between each of these *E. coli* mutations and a set of *C. elegans* mitochondrial mutant strains. Particular *E. coli* mutations that affect the import of the iron siderophore enterobactin into the cytoplasm of *E. coli* or the retrieval of iron in the bacterial cytoplasm from the imported enterobactin bound to iron showed a dramatic synthetic lethality with weak alleles of *C. elegans* mitochondrial electron transport chain components which when grown on wild type bacteria are very healthy. Because Fe(III) is insoluble in aerobic environments, bacteria produce siderophores to retrieve Fe(III). Enterobactin binds Fe(III) with a Km of 10^−39^ M; the enterobactin bound to Fe(III) is then retrieved from outside of the cell or periplasm by specific *E. coli* enterobactin::Fe receptors and the tightly bound Fe(III) is removed from enterobactin in the bacterial cytoplasm by a dedicated enterobactin esterase (5). But *E. coli* mutations that disrupt enterobactin biosynthesis genes were not synthetic lethal with *C. elegans* mitochondrial mutants. In fact, *E. coli* double mutants defective for both enterobactin synthesis and enterobactin import, or for both enterobactin synthesis and Fe(III) removal were no longer synthetic lethal with the *C. elegans* mitochondrial mutants. Thus, the production of enterobactin and a failure to remove the covalently bound iron from enterobactin is required for the toxic interaction between siderophore uptake or esterase mutant *E. coli* and *C. elegans* mitochondrial mutants. Antioxidants such as ascorbic acid or resveratrol also strongly suppressed the inviability of *C. elegans* mitochondrial mutants grown on the *E. coli* enterobactin siderophore utilization or import mutants. We hypothesize that ROS produced by *C. elegans* mitochondrial mutations when combined with ferric-chelated enterobactin to induce increased ROS, which leads to lethality. These data point to ROS as a possible interkingdom communication channel between the microbiome and the microbial-derived endosymbiont of eukaryotes, the mitochondrion.

## Results

To systematically identify bacterial mutations that may affect mitochondrial function, we fed wild type *C. elegans* carrying a mitochondrial unfolded protein response chaperone gene *hsp-6::gfp* with individual bacterial mutant strains from the *E. coli* Keio collection library and screened for *E. coli* mutations that induce the mitochondrial unfolded protein response (UPR^mt^ and/or slow *C. elegans* development. The Keio library is ~4000 single-gene in-frame insertions of a kanamycin resistance cassette in the *E. coli* K12 BW25113 background (6). *C. elegans hsp-6* encodes a mitochondrial matrix HSP70 chaperone that is specifically up-regulated by mutations or drugs that impair mitochondrial function and is a standard marker for the UPR^mt^ (7). We inoculated each of the 4000 Keio collection *E. coli* strains individually on the surface of an agar NGM *C. elegans/E* coli growth agar, added about 100 larval stage one wild type *C. elegans* carrying *hsp-6::GFP* and screened for induction of *hsp-6::GFP* using a dissecting scope fluorescence microscope. We also screened the same wells for slower than normal growth of the *C. elegans* strain. We identified 45 mutant *E. coli* strains that robustly induced *hsp-6::gfp* (Supplementary dataset S1, Fig. S1 and Fig. S2). In the same screening wells observed the next day, we screened the same *E. coli* mutants again for a delay in *C. elegans* rate of development from larval stage one to adulthood. Twenty-four of these 45 *E. coli* mutants also slowed wild type *C. elegans* growth (Table 1). All 24 of the *E. coli* mutants that cause growth delay also were retrieved in the set of 45 *E. coli* mutants that activated *C. elegans hsp-6::gfp*, suggesting that mitochondrial dysfunction is a major axis of bacterial/host nutritional interaction (Table 1). Surprisingly, none of the ~4000 viable *E. coli* Keio mutants that were screened failed to supply some essential micronutrient or macronutrient or produced a toxin that disallowed wild type *C. elegans* growth. Thus, 45 *E. coli* mutations induce a mitochondrial stress response, and half of these also slowed wild type *C. elegans* growth, but none of the Keio mutant collection was missing any essential bacterial factor (such as heme) for *C. elegans* growth. Our screen would have easily detected such a lethal interaction. There are 303 *E. coli* Keio gene disruptions that are inviable and thus not easily tested for *C. elegans* interactions.

**Table 1:**
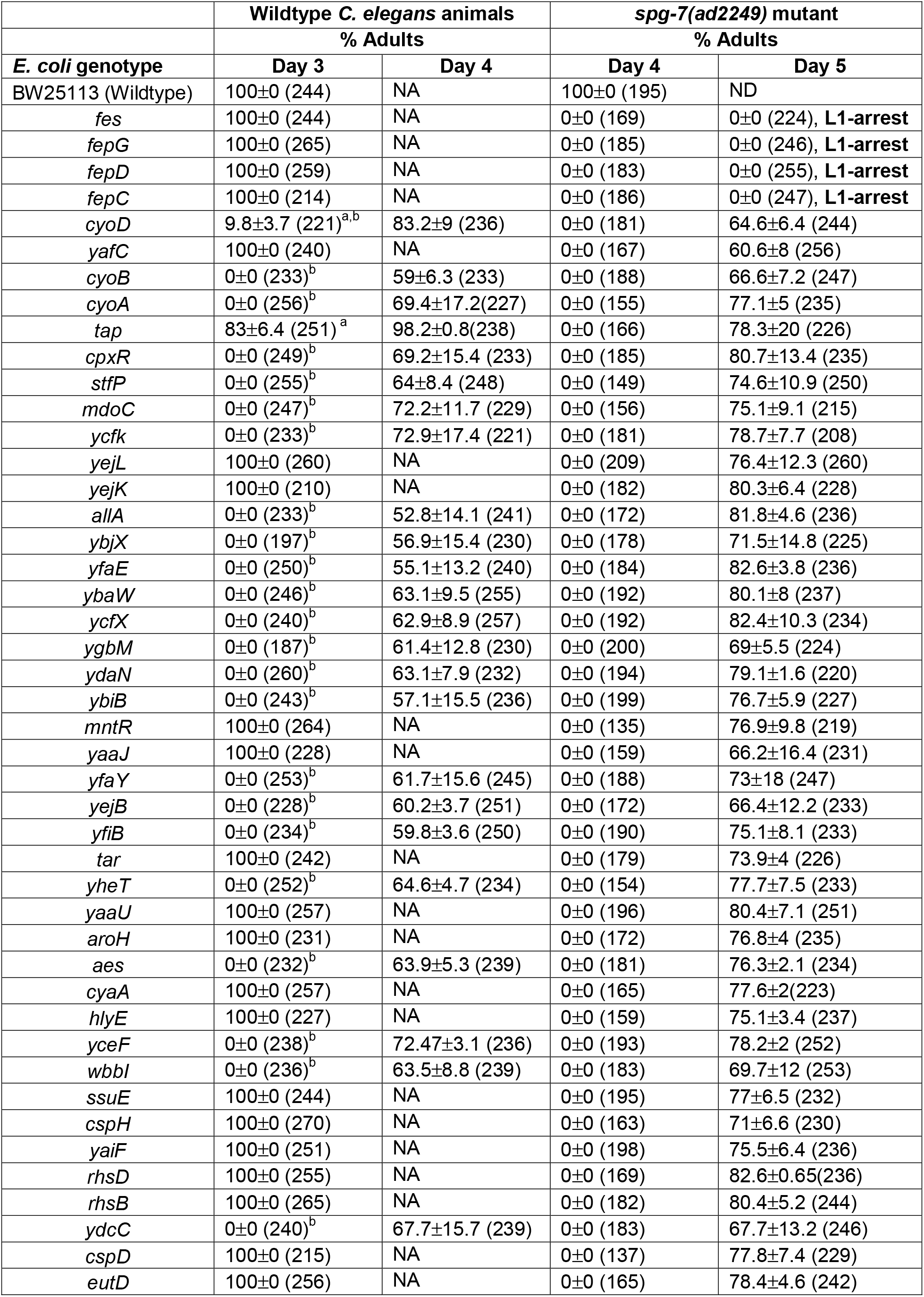
Quantification of developmental delay phenotype in wildtype background and *spg-7(ad2249)* mutant background. Synchronized L1 larval stage wildtype animal were inoculated onto NGM media plates seeded with individual Keio *E. coli* mutant strains and incubated at 20°C. The number of adult animals and the total number of animals were counted on day 3 and day 4 of feeding wildtype animals on Keio *E. coli* mutant strains. Data from three independent trials were collected; the average of percentage of adults and the standard deviation are shown. The total number of animals counted in all the three independent trials is shown in the parentheses. ^a^Compared to Wildtype animals fed on BW25113 on day 3, Unpaired t-test, P<0.0001. ^b^Since there were no adults or only few adults on these plates, the plates were scored on day 4 for the presence of adults. While there were no adults on day3, many of the wells had animals that reached adulthood on day 4. NA, Not applicable since all the animals were adults on day 3 itself. For column 4 and 5, synchronized L1 larval stage *spg-7(ad2249)* mutant were inoculated onto NGM media plates seeded with individual Keio *E. coli* mutant strains and incubated at 20°C. The number of adult animals and the total number of animals were counted on day 4 and 5 of feeding *spg-7(ad2249* animals on Keio *E. coli* mutant strains. Data from three independent trials were collected; the average of percentage of adults and the standard deviation are shown. The total number of animals counted in all the three independent trials is shown in the parentheses.

|  | Wildtype *C. elegans* animals |  | *spg-7(ad2249)* mutant |  |
| --- | --- | --- | --- | --- |
|  | % Adults |  | % Adults |  |
| *E. coli* genotype | Day 3 | Day 4 | Day 4 | Day 5 |
| BW25113 (Wildtype) | 100±0 (244) | NA | 100±0 (195) | ND |
| *fes* | 100±0 (244) | NA | 0±0 (169) | 0±0 (224), **L1-arrest** |
| *fepG* | 100±0 (265) | NA | 0±0 (185) | 0±0 (246), **L1-arrest** |
| *fepD* | 100±0 (259) | NA | 0±0 (183) | 0±0 (255), **L1-arrest** |
| *fepC* | 100±0 (214) | NA | 0±0 (186) | 0±0 (247), **L1-arrest** |
| *cyoD* | 9.8±3.7 (221)<sup>a,b</sup> | 83.2±9 (236) | 0±0 (181) | 64.6±6.4 (244) |
| *yafC* | 100±0 (240) | NA | 0±0 (167) | 60.6±8 (256) |
| *cyoB* | 0±0 (233)<sup>b</sup> | 59±6.3 (233) | 0±0 (188) | 66.6±7.2 (247) |
| *cyoA* | 0±0 (256)<sup>b</sup> | 69.4±17.2(227) | 0±0 (155) | 77.1±5 (235) |
| *tap* | 83±6.4 (251)<sup>a</sup> | 98.2±0.8(238) | 0±0 (166) | 78.3±20 (226) |
| *cpxR* | 0±0 (249)<sup>b</sup> | 69.2±15.4 (233) | 0±0 (185) | 80.7±13.4 (235) |
| *stfP* | 0±0 (255)<sup>b</sup> | 64±8.4 (248) | 0±0 (149) | 74.6±10.9 (250) |
| *mdoC* | 0±0 (247)<sup>b</sup> | 72.2±11.7 (229) | 0±0 (156) | 75.1±9.1 (215) |
| *ycfk* | 0±0 (233)<sup>b</sup> | 72.9±17.4 (221) | 0±0 (181) | 78.7±7.7 (208) |
| *yejL* | 100±0 (260) | NA | 0±0 (209) | 76.4±12.3 (260) |
| *yejK* | 100±0 (210) | NA | 0±0 (182) | 80.3±6.4 (228) |
| *allA* | 0±0 (233)<sup>b</sup> | 52.8±14.1 (241) | 0±0 (172) | 81.8±4.6 (236) |
| *ybjX* | 0±0 (197)<sup>b</sup> | 56.9±15.4 (230) | 0±0 (178) | 71.5±14.8 (225) |
| *yfaE* | 0±0 (250)<sup>b</sup> | 55.1±13.2 (240) | 0±0 (184) | 82.6±3.8 (236) |
| *ybaW* | 0±0 (246)<sup>b</sup> | 63.1±9.5 (255) | 0±0 (192) | 80.1±8 (237) |
| *ycfX* | 0±0 (240)<sup>b</sup> | 62.9±8.9 (257) | 0±0 (192) | 82.4±10.3 (234) |
| *ygbM* | 0±0 (187)<sup>b</sup> | 61.4±12.8 (230) | 0±0 (200) | 69±5.5 (224) |
| *ydaN* | 0±0 (260)<sup>b</sup> | 63.1±7.9 (232) | 0±0 (194) | 79.1±1.6 (220) |
| *ybiB* | 0±0 (243)<sup>b</sup> | 57.1±15.5 (236) | 0±0 (199) | 76.7±5.9 (227) |
| *mntR* | 100±0 (264) | NA | 0±0 (135) | 76.9±9.8 (219) |
| *yaaJ* | 100±0 (228) | NA | 0±0 (159) | 66.2±16.4 (231) |
| *yfaY* | 0±0 (253)<sup>b</sup> | 61.7±15.6 (245) | 0±0 (188) | 73±18 (247) |
| *yejB* | 0±0 (228)<sup>b</sup> | 60.2±3.7 (251) | 0±0 (172) | 66.4±12.2 (233) |
| *yfiB* | 0±0 (234)<sup>b</sup> | 59.8±3.6 (250) | 0±0 (190) | 75.1±8.1 (233) |
| *tar* | 100±0 (242) | NA | 0±0 (179) | 73.9±4 (226) |
| *yheT* | 0±0 (252)<sup>b</sup> | 64.6±4.7 (234) | 0±0 (154) | 77.7±7.5 (233) |
| *yaaU* | 100±0 (257) | NA | 0±0 (196) | 80.4±7.1 (251) |
| *aroH* | 100±0 (231) | NA | 0±0 (172) | 76.8±4 (235) |
| *aes* | 0±0 (232)<sup>b</sup> | 63.9±5.3 (239) | 0±0 (181) | 76.3±2.1 (234) |
| *cyaA* | 100±0 (257) | NA | 0±0 (165) | 77.6±2(223) |
| *hlyE* | 100±0 (227) | NA | 0±0 (159) | 75.1±3.4 (237) |
| *yceF* | 0±0 (238)<sup>b</sup> | 72.47±3.1 (236) | 0±0 (193) | 78.2±2 (252) |
| *wbbI* | 0±0 (236)<sup>b</sup> | 63.5±8.8 (239) | 0±0 (183) | 69.7±12 (253) |
| *ssuE* | 100±0 (244) | NA | 0±0 (195) | 77±6.5 (232) |
| *cspH* | 100±0 (270) | NA | 0±0 (163) | 71±6.6 (230) |
| *yaiF* | 100±0 (251) | NA | 0±0 (198) | 75.5±6.4 (236) |
| *rhsD* | 100±0 (255) | NA | 0±0 (169) | 82.6±0.65(236) |
| *rhsB* | 100±0 (265) | NA | 0±0 (182) | 80.4±5.2 (244) |
| *ydcC* | 0±0 (240)<sup>b</sup> | 67.7±15.7 (239) | 0±0 (183) | 67.7±13.2 (246) |
| *cspD* | 100±0 (215) | NA | 0±0 (137) | 77.8±7.4 (229) |
| *eutD* | 100±0 (256) | NA | 0±0 (165) | 78.4±4.6 (242) |

To verify the UPR^mt^ induction, we showed that the 45 mutant *E. coli* strains that activate *hsp-6::gfp*, also activate *hsp-60::gfp*, another marker for the UPR^mt^ (Supplementary dataset S1). The *C. elegans* stress response induced by the 45 *E. coli* Keio mutants is specific for mitochondria because none of the 45 bacterial mutants caused activation of *hsp-4::gfp* (Supplementary dataset S1), a reporter of the unfolded protein response in the endoplasmic reticulum (UPR^er^) (8). Similarly, the 45 *E. coli* Keio mutants did not activate *pgp-5::gfp* (Supplementary dataset S1), a reporter of a translational stress response (9). ATFS-1 is a transcription factor required for the activation of UPR^mt^ genes, such as *hsp-6* and *hsp-60*. We tested each of the Keio *E. coli* mutants on a *C. elegans atfs-1(tm4525); hsp-60::GFP C. elegans* strain and found that all 45 bacterial mutants failed to induce the *hsp-60::GFP* UPR^mt^ in the *atfs-1(tm4525)* background (Supplementary dataset S1). Thus, these *E. coli* mutants activate the *C. elegans* UPRmt via the expected ATFS-1 transcription factor.

Because all of the *E. coli* mutants that slowed growth induce the mitochondrial-unfolded response, and some are implicated in the *E. coli* electron transport chain, we examined whether any of these *E. coli* mutants strongly interact with *C. elegans* mutations in nuclear-encoded mitochondrial proteins. Previous *C. elegans* RNAi screens have shown that many mitochondrial gene inactivations are lethal (10). However, viable reduction-of-function mutants have been characterized (11). We tested a viable *spg-7(ad2249)* mutant (12). *spg-7* encodes the human homolog of AFG3L2, a conserved m-AAA metalloprotease that processes particular proteins and protein complexes in the mitochondria (12). Although feeding many of the 45 Keio collection *E. coli* mutants caused developmental delay in *spg-7(ad2249)*, four of the 45 Keio mutants caused a dramatic arrest phenotype in *spg-7(ad2249)*: *Δfes::kan*, *ΔfepD::kan*, *ΔfepG::kan*, *ΔfepC::kan* mutants (Table 1). For example, *Δfes::kan* feeding induces a highly penetrant very dramatic developmental arrest with *C. elegans spg-7(ad2249)* while wildtype *C. elegans* fed on *Δfes::kan* are grow normally (Fig. 1A). Based on the morphological features, animal size measurements, and the germline development stage, *Δfes::kan* feeding in *spg-7(ad2249)* mutant causes an larval stage one (L1) arrest (Fig. S3A-C). Each of these *E. coli* genes that cause a synthetic arrest with C. elegans mitochondrial mutant strains mediates steps in the retrieval of iron from the *E. coli* siderophore enterobactin. Enterobactin is a high-affinity iron siderophore produced by many gram-negative bacteria, including *E. coli* (5, 13). *E. coli* produce enterobactin under iron deficiency via the proteins encoded by the *entCDEBAH* operon (Fig. 1B-C). Enterobactin is secreted into the environment, the soil or in a cell, where it binds to iron, forming a complex which is imported into *E. coli* through the transporters located in the bacterial outer membrane (13) (Fig. 1C). The ferric-enterobactin complex is transported into the cytoplasm via the *fepDGC* ATP-binding cassette transporter (14, 15). FepB encodes a periplasmic protein that binds ferric enterobactin and directs it to inner membrane transporters, FepD and FepG, which pump it into the bacterial cytoplasm. FepC assists FepD and FepG in transporting ferric enterobactin from the periplasm into the cytoplasm (Fig. 1C) (16). Fes encodes enterobactin esterase that catalyzes the hydrolysis of both enterobactin and ferric enterobactin. In the *E. coli* cytoplasm, Fes hydrolyses enterobactin to release iron and linearized 2,3-dihydroxy-N-benzoyl-L-serine trimer as well as the dimer and monomer (Fig. 1C) (17–19). Genetic disruption of E. coli *fes* causes accumulation of the unhydrolyzed ferric enterobactin in the bacterial cytoplasm (20, 21) whereas genetic disruption fepDGC causes accumulation of the ferric enterobactin in the *E. coli* periplasm where the cytoplasmic Fes esterase cannot release iron from its nearly covalent bond in ferric enterobactin (14, 15) (Fig. S4A). This complex enterobactin synthesis and import biology is not parochial to *E. coli:* siderophores related to enterobactin are used across bacterial phylogeny, and the hijacking of iron-loaded siderophores produced by one species and taken up by another bacterial species is a key feature of the iron competition landscape.

**Figure 1:**
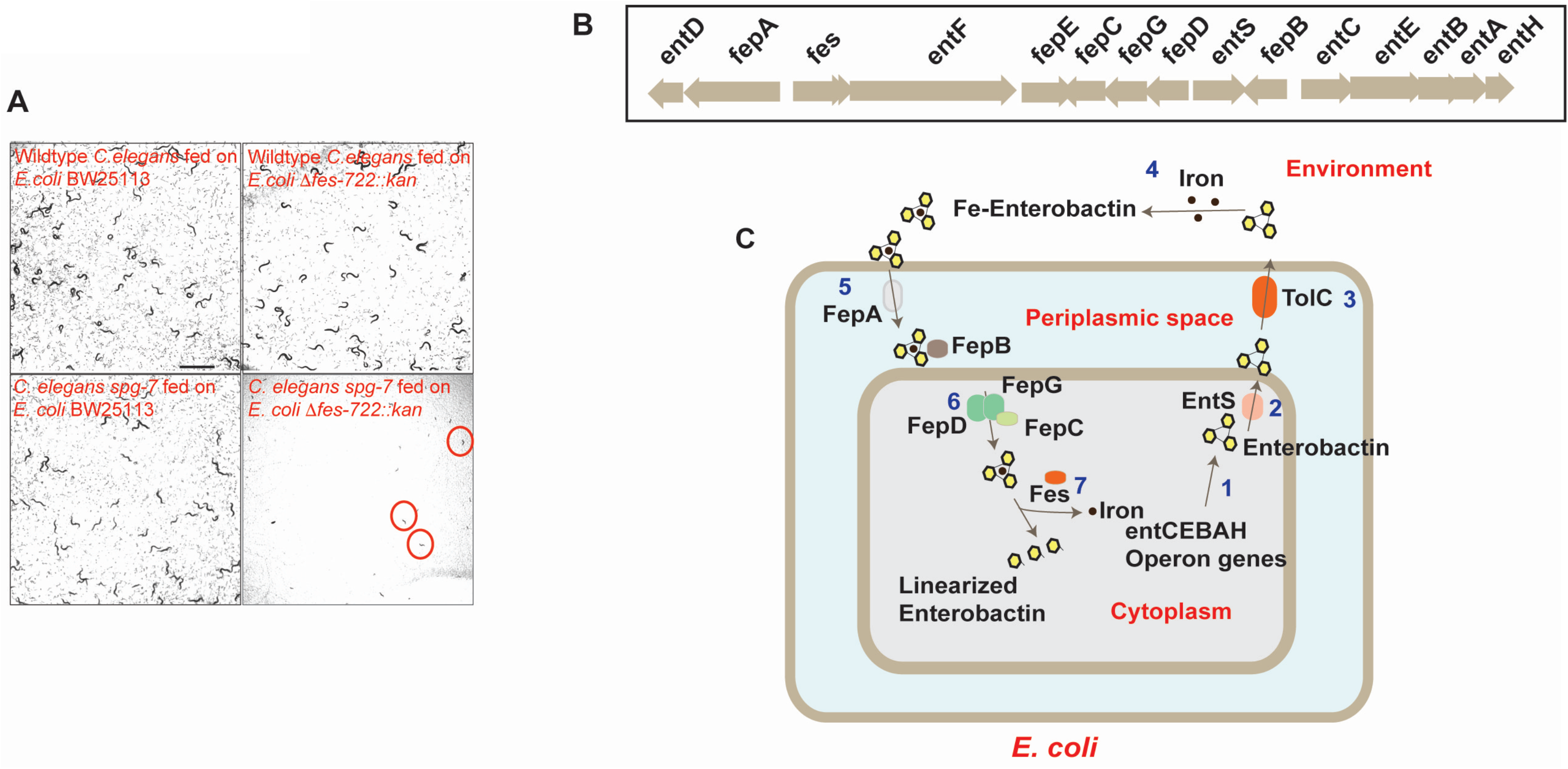
*E. coli* mutations that affect enterobactin siderophore utilization or import are synthetic lethal with *C. elegans* mitochondrial mutations. A) *spg-7(ad2249)* animals grown on *Δfes::kan E. coli* arrest as L1-larvae. Scale bar, 1000μM. B) Operon structure of genes in the enterobactin pathway C) Diagrammatic representation of enterobactin biosynthetic pathway in *E. coli*.

To determine whether other *E. coli* genes required for enterobactin production cause the dramatic L1-synthetic larval arrest in the *spg-7 C. elegans* mitochondrial mutant, we tested each of the *E. coli* enterobactin pathway gene mutants on the *C. elegans spg-7* mutant. Feeding *C. elegans spg-7(ad2249)* mutants on *ΔentC::kan*, *ΔentD::kan*, *ΔentE::kan*, *ΔentB::kan*, *ΔentA::kan*, *ΔentH::kan*, which are defective in the production of enterobactin, does not cause L1-larval arrest (Table 2). This suggests that it is not lack of enterobactin or *E. coli* iron starvation that causes developmental arrest in the *C. elegans spg-7* mitochondrial mutant but it is the accumulation of iron-chelated enterobactin in the cytoplasm or periplasm of *E. coli* or iron-chelated enterobactin that is transported from the *E. coli* to the *C. elegans* cells with electron transport mutations that causes arrest of the *C. elegans* mitochondrial mutant.

**Table 2:** Feeding Ferric-Enterobactin induces developmental arrest in *C. elegans* mitochondrial mutants

| Fed on | Description | % Larval arrest (N) |  |
| --- | --- | --- | --- |
|  |  | Wildtype | *spg-7(ad2249)* |
| BW25113 | Wildtype | 0±0 (179) | 0±0 (219) |
| *ΔfepC726::kan* | Ferric enterobactin transport; ATP-binding protein *fepC* | 0±0 (132) | 100±0 (178) |
| *ΔfepG727::kan* | Ferric enterobactin transport system permease protein *FepG* | 0±0 (195) | 100±0 (196) |
| *ΔfepD728::kan* | Ferric enterobactin transport system permease protein *FepD* | 0±0 (198) | 100±0 (181) |
| *ΔfepB730::kan* | Ferrienterobactin-binding periplasmic protein | 0±0 (231) | 0±0 (174) |
| *ΔfepA721::kan* | Ferrienterobactin receptor | 0±0 (159) | 0±0 (188) |
| *Δfiu-777::kan* | Catecholate siderophore receptor *Fiu* | 0±0 (188) | 0±0 (164) |
| *ΔcirA782::kan* | Colicin I receptor; Postulated to participate in iron transport | 0±0 (201) | 0±0 (190) |
| *Δfes-722::kan* | Enterochelin esterase | 0±0 (151) | 100±0 (205) |
| *ΔentF724::kan* | Enterobactin synthase component *F* | 0±0 (142) | 0±0 (165) |
| *ΔentC731::kan* | Isochorismate synthase *EntC* | 0±0 (166) | 0±0 (179) |
| *ΔentE732::kan* | Enterobactin synthase component *E* | 0±0 (202) | 0±0 (173) |
| *ΔentB733::kan* | Enterobactin synthase component *B* | 0±0 (191) | 0±0 (172) |
| *ΔentA734::kan* | 2,3-dihydro-2,3-dihydroxybenzoate dehydrogenase | 0±0 (183) | 0±0 (162) |

Synchronized L1 larval stage wildtype or *spg-7(ad2249)* mutant were inoculated onto NGM media plates seeded with individual Keio *E. coli* mutant strains and incubated at 20°C. The number of L1-arrest animals and the total number of animals were counted on day 3.

To determine whether the L1 larval arrest caused by feeding *Δfes::kan* mutant is specific for *spg-7(ad2249)* mutant or a more generalized phenotype of many *C. elegans* mitochondrial mutants, we tested other viable reduction-of-function *C. elegans* mitochondrial mutants (Table 3). *C. elegans nduf-7(et19), isp-1(qm150)*, and *clk-1(qm30)* mutant animals grown on the *E. coli Δfes::kan* mutant also dramatically arrest as L1 larvae (Table 3; Fig. S5) while particular other *C. elegans* mitochondrial mutants including *gas-1(fc21), nduf-2.2(ok2397), nuo-6(qm200), mev-1(kn1)*, and *ucr-2.3(pk732)* did not arrest when grown on the *E. coli Δfes::kan* mutant (Table 3; Fig. S5). *nduf-7(et19)* is a partial loss-of-function mutation in *nduf-7* (NADH-ubiquinone oxidoreductase Fe-S), a subunit of the mitochondrial electron transport chain complex I (11, 22). *nduf-7(et19)* mutant animals strongly activate UPR^mt^, have a reduced respiration rate and longer lifespan (22). *isp-1* encodes a Rieske iron sulphur protein (ISP) that functions in the cytochrome b-c1 complex III subunit of the mitochondrial respiratory chain (11). *isp-1(qm150)* mutants have lower oxygen consumption, decreased ROS production, and increased lifespan (23). *clk-1* encodes an ortholog of the highly conserved demethoxyubiquinone (DMQ) hydroxylase that is necessary for ubiquinone biosynthesis (11). Mutations in *C. elegans clk-1* causes slow development, reduced respiration, and increased lifespan when grown on an *E. coli* strain that is competent to produce bacterial ubiquinone that is identical except for a shorter isoprenoid chain that localizes this electron carrier to the membrane (24).

**Table 3:**
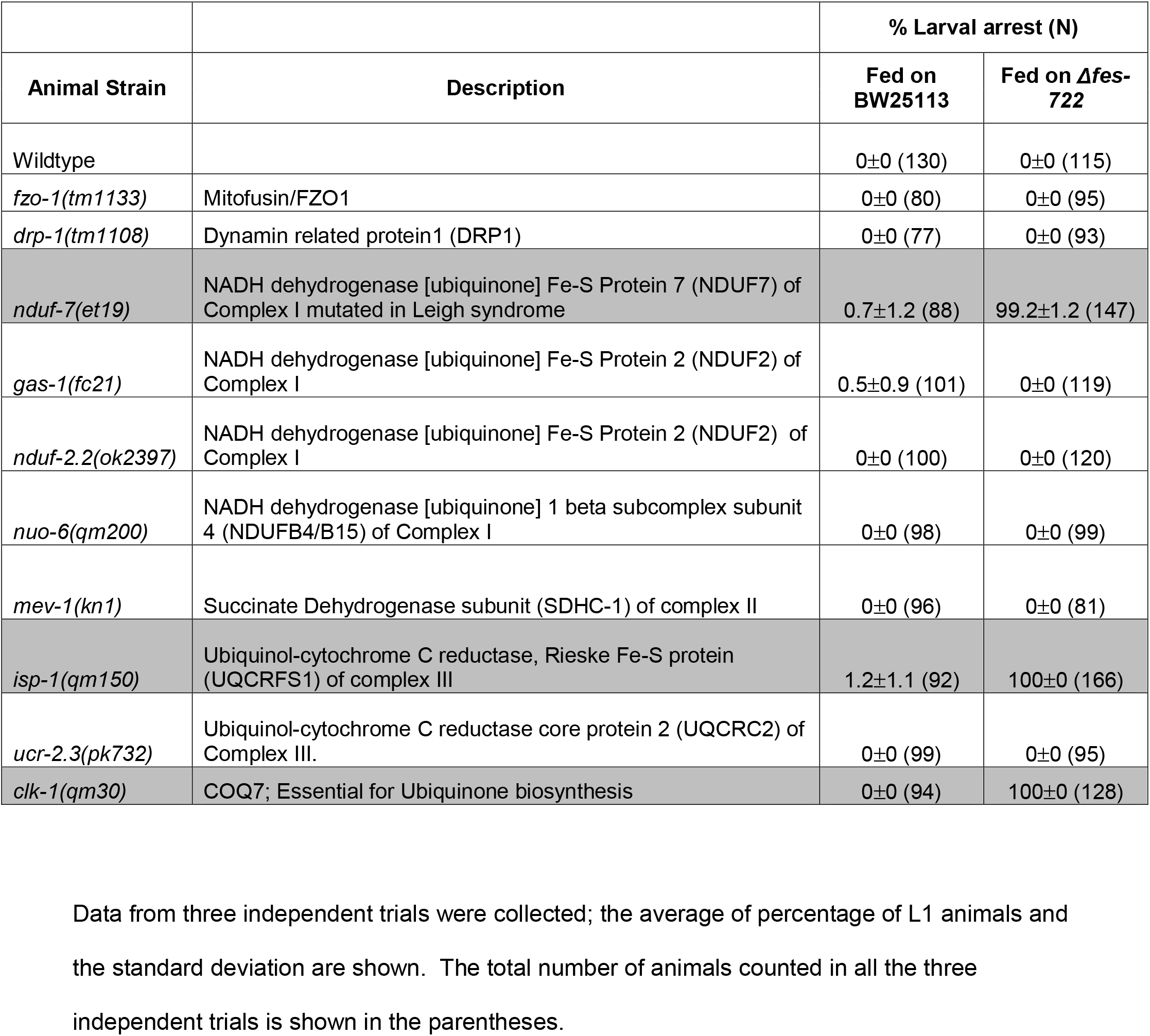
Table showing list of the mitochondrial mutants tested to potential interaction with *Δfes* mutant

Because *spg-7, nduf-7, isp-1*, and *clk-1* mediate distinct steps in mitochondrial respiratory pathways, the arrest induced by feeding mitochondrial mutants on *E. coli Δfes::kan* mutant is likely due to a toxic interaction between the mitochondrial homeostasis induction (one aspect of which is *hsp-6* expression) induced by the *spg-7, nduf-7, isp-1*, and *clk-1* mutations (but not by the *gas-1(fc21), nduf-2.2(ok2397), nuo-6(qm200), mev-1(kn1)*, and *ucr-2.3(pk732)* mutations) and the iron toxicity caused by *E. coli* enterobactin mutants (Table 3).

Why do mutations that affect enterobactin-iron uptake or utilization but not enterobactin production cause developmental arrest in combination with particular *C. elegans* mitochondrial mutations? Our hypothesis is that the *E. coli* mutants that carry ferric enterobactin, but not *E. coli* mutants that make no enterobactin, are toxic for animals with the *spg-7, nduf-7, isp-1*, and *clk-1* mitochondrial dysfunctions. Inactivation of *fes* causes accumulation of the unhydrolyzed ferric enterobactin in the *E. coli* cytoplasm (20, 21) whereas inactivation in FepDGC causes accumulation of the ferric enterobactin in the *E. coli* periplasm (14, 15) which is inaccessible for cytoplasmic Fes to act upon (Fig. S4A). As *C. elegans* consumes the Fes or Fep mutant *E. coli*, the ferric enterobactin may be unaltered by *C. elegans* esterases, which may not recognize this nearly covalent iron chelation. This ferric enterobactin may be transported into *C. elegans* cells, where if mitochondria are defective, perhaps producing ROS (see Discussion), this ferric enterobactin is now toxic.

*fes* is in an operon with *ybdZ* and *entF*, which are also involved in enterobactin biosynthesis. Because bacterial insertion drug resistant cassette mutations in one gene in the operon are often polar on downstream genes, we excised the Keio collection antibiotic resistance cassette insertion in *Δfes::kan* to create an in-frame non-polar deletion allele of the single *fes* gene in the operon, which we call *Δfes*. Feeding *spg-7(ad2249)* mutant animals on *Δfes E. coli* arrest also caused developmental arrest, suggesting that that the arrest phenotype is due to the absence of Fes protein, not polar effects on downstream genes (Fig. 2A-B). To test whether the developmental arrest of the *C. elegans* mitochondrial mutants is due to ferric enterobactin, we inactivated enterobactin biosynthesis in the *Δfes* mutant background by constructing *Δfes entA::kan* and *Δfes entB::kan* double mutants. While 100% of *C. elegans spg-7(ad2249)* mutants arrest when grown on *Δfes* mutant *E. coli*, 0% of *spg-7(ad2249)* mutants arrest when grown the *Δfes entA::kan* or *Δfes entB::kan* double mutant *E. coli* that does not generate enterobactin and therefore does not generate ferric enterobactin (Fig. S4B; Fig. 2A-B). Further, to test whether ferric enterobactin is required within *E. coli* to produce the arrest in *spg-7(ad2249)* mutants, we constructed *Δfes cirA782::kan, Δfes fepA::kan*, and *Δfes fiu::kan* double mutants (Fig. 2B). FepA is an outer membrane protein that binds and transports ferric enterobactin into the periplasm of *E. coli* (25, 26). Fiu is an outer membrane protein that mediates uptake of dihydroxybenzoylserine, a breakdown product of enterobactin (27). Cir is another outer membrane protein that mediates uptake of ferric-enterobactin and other breakdown products of enterobactin (27). Although FepA, Fiu and CirA mediate import of ferric enterobactin into the bacterial periplasm, FepA is the major transporter necessary for the uptake of ferric enterobactin into bacterial periplasm. While 100% of *C. elegans spg-7(ad2249)* mutants arrest when grown on *Δfes E. coli, Δfes cirA::kan* or *Δfes fiu::kan* double mutant *E. coli*, 0% of *spg-7(ad2249)* mutants arrest in the *Δfes fepA::kan* double mutant *E. coli* (Fig. 2B). This result suggests that blocking uptake of ferric enterobactin from the environment into the *E. coli* is sufficient to suppress the L1-larval arrest of *C. elegans* mitochondrial mutants grown on *Δfes E. coli* mutant.

**Figure 2:**
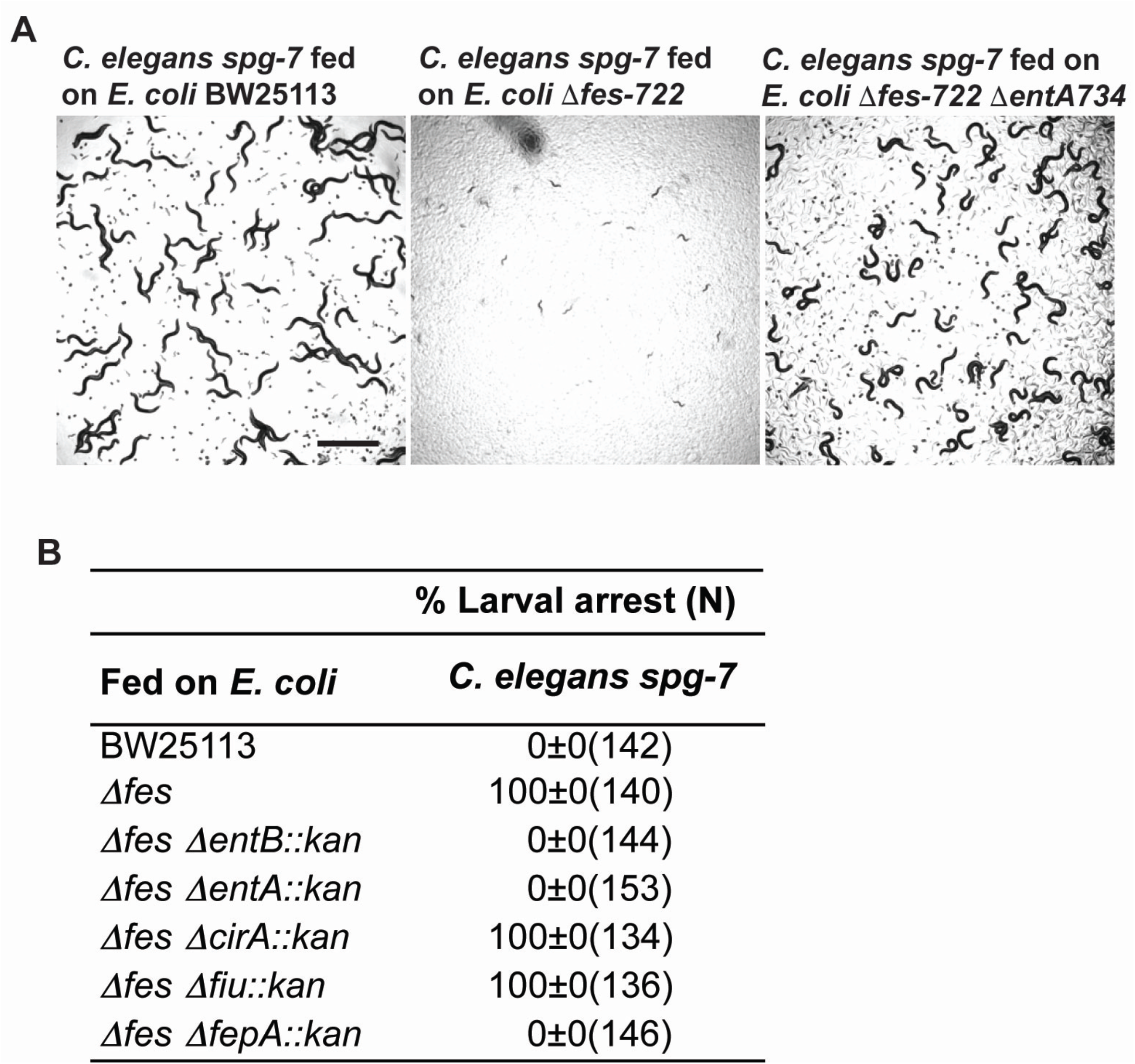
Enterobactin is necessary for the developmental arrest phenotype of *C. elegans* mitochondrial mutants. A) *spg-7(ad2249)* animals grown on wildtype *E.coli* BW25113 develop normally while *spg-7(ad2249)* mutant grown in *Δfes* mutant arrest as L1-larvae. In contrast, *spg-7(ad2249)* animals grown on *Δfes ΔentA::kan* double mutants develop normally. Scale bar, 1000μM. B) Table showing suppression of the larval arrest phenotype in animals grown on *Δfes ΔentA::kan, Δfes ΔentB::kan*, and *Δfes ΔfepA::kan* mutants. Data from three independent trials were collected; the average of percentage of adults and the standard deviation are shown. The total number of animals counted in all the three independent trials is shown in the parentheses.

Because the *E. coli* enterobactin uptake and iron retrieval mutants activate a mitochondrial unfolded protein response even in a wild type *C. elegans* that grows at the normal rate (Table 1), we assessed the effect of these *E. coli* mutants on *C. elegans* mitochondrial morphology. The mitochondria of wildtype animals grown on wild type *E. coli* BW25113 have tubular or slightly elongated morphology, but the mitochondria in the hypodermis of wildtype animals grown on *Δfes E. coli* are fragmented as assessed by Nonyl Acridine Orange (NAO), a membrane potential-independent mitochondrial dye (Fig. 3A). We also used a transgenic strain that expresses mitochondrial outer membrane protein TOMM-20 tagged with mRFP in the body wall muscles of *C. elegans*. The mitochondrial morphology is tubular and elongated in animals grown on wild type *E. coli* BW25113 whereas wild type *C. elegans* grown on *Δfes E. coli* show a more fragmented mitochondria in the body wall muscle (Fig. 3B). Consistent with the abnormal mitochondrial morphology, ATP levels are significantly reduced in wild type *C. elegans* grown on *Δfes E. coli* compared to wild type *E. coli* BW25113 (Fig. 3C-D). These data show that the induction of the *hsp-6* in wild type *C. elegans* grown on the *E. coli* iron-siderophore retrieval mutants accurately reports their mitochondrial defects.

**Figure 3:**
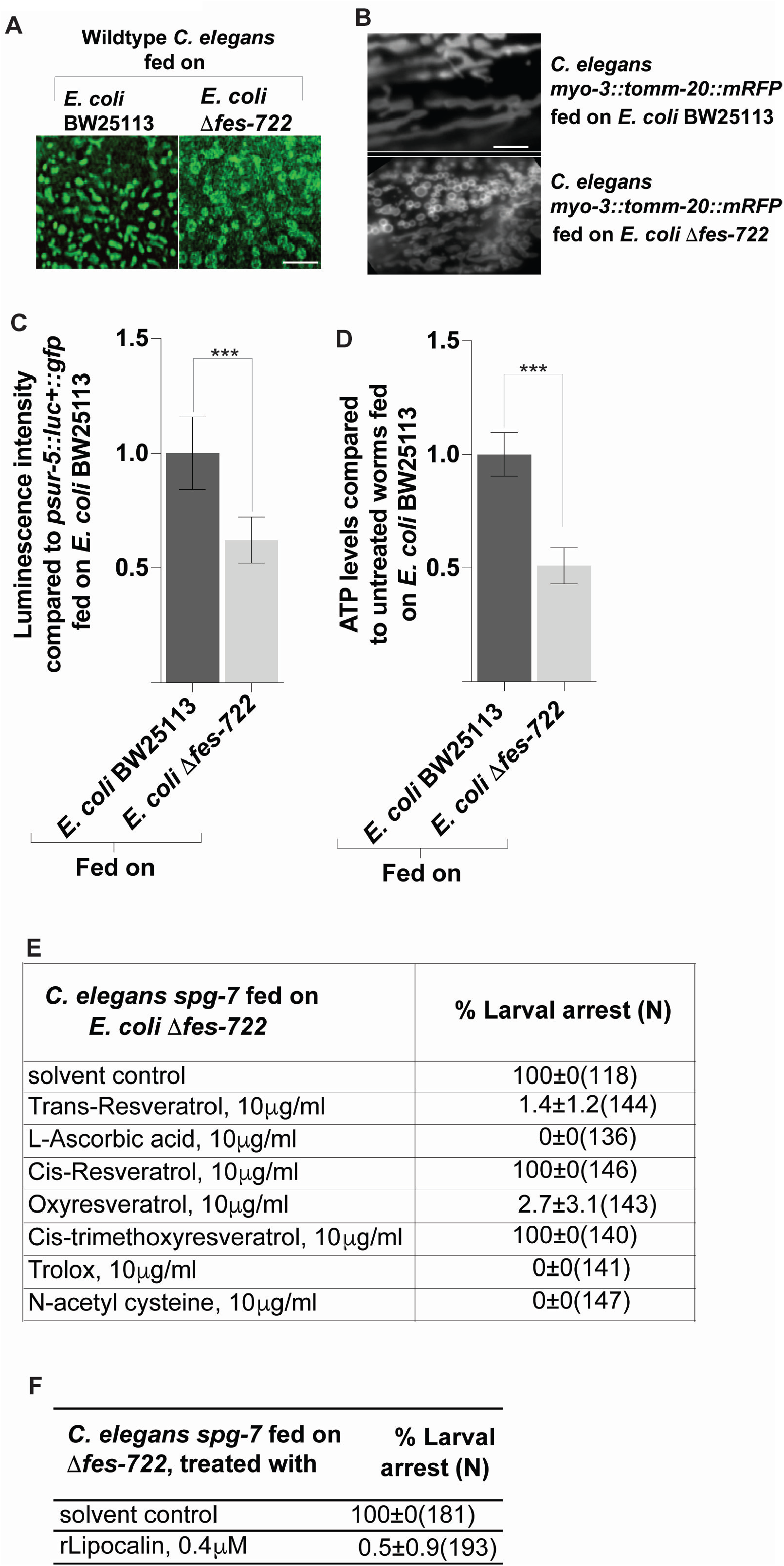
Mitochondrial structure and function are disrupted in wildtype animals grown on Δ*fes* mutant *E. coli*. A) Wildtype animals grown on *Δfes* mutant *E. coli* display fragmented hypodermal mitochondrial morphology as assessed using NAO. Scale bar, 100μM. B) Transgenic animals expressing TOMM-20::mRFP in the muscles grown on *Δfes* mutant *E. coli* display a fragmented mitochondrial morphology. Scale bar, 100μM. C) Animals grown on *Δfes* mutant *E. coli* display lowered ATP levels compared to animals grown on *E.coli* BW25113 as assessed using a strain that expresses luciferase in all tissues. Unpaired t-test; *** P< 0.001. Mean ± s.d of n=10. D) ATP levels in animals grown on *Δfes* mutant *E. coli* display lowered ATP levels compared to animals grown on *E.coli* BW25113. Unpaired t-test; *** P< 0.001. Mean ± s.d of n=3. E) While antioxidant treatments suppress the L1-larval arrest phenotype induced by feeding *Δfes* mutants to *C. elegans* mitochondrial mutants, cis-resveratrol and cis-trimethoxyresveratrol fails to suppress the arrest phenotype. While trans-resveratrol and oxyresveratrol suppressed arrest, cis-resveratrol or cis-trimethoxy resveratrol did not suppress the arrest: perhaps these isomers distinguish between the free radical production surface. Data from three independent trials were collected; the average of percentage of adults and the standard deviation are shown. The total number of animals counted in all the three independent trials is shown in the parentheses. F) Mouse lipocalin is sufficient to suppress the larval arrest phenotype of *spg-7(ad2249)* animals grown on *Δfes* mutants. Data from three independent trials were collected; the average of percentage of adults and the standard deviation are shown. The total number of animals counted in all the three independent trials is shown in the parentheses.

*spg-7(ad2249)* animals grown on gentamicin-treated or tetracycline-treated *E. coli Δfes* arrested at L1-arrest stage, showing that even non-growing bacteria could supply the toxic iron-siderophore product to a *C. elegans* mitochondrial mutant (Table S1). But ampicillin-killed *E. coli Δfes* did not arrest with *C. elegans spg-7(ad2249)* (Table S1). Tetracycline, kanamycin, and gentamicin are bacteriostatic whereas ampicillin targets cell wall biosynthesis and is bactericidal. Ampicillin may lyse the bacterial cell to release ferric enterobactin before the bacterial cell is engulfed in the *C. elegans* intestine to absorb ferric enterobactin.

To determine the relative dose of *E. coli Δfes* needed for this toxic interaction, we mixed various proportions of *E. coli Δfes* and wildtype BW25113. *C. elegans spg-7(ad2249)* grown on 50% live wildtype *E. coli* BW25113 and 50% live *E. coli Δfes::kan* did not arrest (Table S1). *spg-7(ad2249)* grown on 50% kanamycin-treated wildtype *E. coli* BW25113 and 50% live *E. coli Δfes::kan*, arrested as L1-larvae (Table S1). A mixture of 75% kanamycin-treated wildtype *E. coli* BW25113 and 25% live *E. coli Δfes* showed the L1-arrest (Table S1), showing that living wildtype *E. coli* BW25113 is required for the suppression of *E. coli Δfes* feeding-induced L1-arrest phenotype. Feeding *C. elegans spg-7(ad2249)* mutant animals on mixture of 1-part live *E. coli* BW25113 in 10,000 parts live *E. coli Δfes* fully suppressed the *E. coli Δfes* arrest; however, killed wildtype *E. coli* could not suppress arrest (Table S1). Differential growth rates of wild type and *fes* mutant *E. coli* on *C. elegans* growth plates explains the potency of the interaction; after 48 hours on NGM plates the 1:10,000 dilution of *E. coli* BW25113 in *E. coli Δfes* had decreased to ~1.3:1 (Table S2). Thus, wildtype *E. coli* outgrows the *Δfes* mutants.

Mutations in *isp-1, nduf-7*, and *spg-7* cause increased ROS production by the mitochondria (12, 22, 28). Elevated ROS levels is essential for the resistance to hemiasterlin, a mitochondrial poison, of the *spg-7(ad2249)* mutant because this is suppressible by NAC (12). We found that *spg-7(ad2249)* animals that were grown on *Δfes E. coli* mutant, but mixed with antioxidants such as ascorbic acid, N-acetyl cysteine (NAC), resveratrol, or trolox completely suppressed the arrest caused by *E. coli* fes mutant fed to *C. elegans spg-7* (Fig. 3E). The fact that several other antioxidants were able to suppress arrest suggests that ROS is the likely cause of the toxic interaction between bacterial mutants that accumulate ferric enterobactin and the *C. elegans* mitochondrial mutants. Elevated ROS in *isp-1(qm150)* contributes to its increased longevity because the longevity increase is suppressed by NAC (28). UPR^mt^ activation in the *nduf-7(et19)* is also suppressed by NAC (22). Consistent with this observation, we found that the L1-larval arrest induced by feeding *Δfes E. coli* to the *nduf-7(et19)* mutant is suppressible by NAC (Table S3). When we fed *C. elegans spg-7(ad2249)* animals the *E. coli Δfes* pregrown in the presence of trans-resveratrol and washed off the trans-resveratrol, the animals arrest, suggesting that the site-of-action of the antioxidants is likely within the animals (Table S4).

Because hydrogen peroxide is one of the major forms of endogenous ROS, we monitored *in vivo* hydrogen peroxide levels of the interaction between the *E. coli Δfes* mutations and *C. elegans*, using HyPer, a peroxide-specific sensor protein. Transgenic wild type animals that express the HyPer sensor (*jrIs1*) (29) were grown either on *E. coli* BW25113 or *E. coli Δfes* and peroxide levels were measured. Endogenous peroxide levels were not significantly different between wild type *C. elegans* grown *E. coli* BW25113 or *E. coli Δfes;* however, addition of exogenous peroxide results in significantly higher peroxide levels in animals grown on *E. coli Δfes* compared to animals grown on *E. coli* BW25113 (Fig. S6). We hypothesize that the exogenous peroxide interacts with ferric-enterobactin and via the Fenton reaction to generate more peroxide.

In the evolutionary “tug-of-war” for iron, animals rely on the enterobactin-binding protein Lipocalin-2 to counteract iron loss to microbes that secrete siderophores (30). Secreted lipocalin 2 sequesters the ferric-enterobactin to limit bacterial growth (30). We tested whether exogenous lipocalin-2 can suppress the arrest of *spg-7(ad2249)* grown on the *E. coli Δfes* mutant. We found that *spg-7(ad2249)* animals that were grown on the *E. coli Δfes* in the presence of 0.4μM of recombinant Mouse lipocalin-2 no longer arrest as L1-Larvae (Fig. 3F).

## Discussion

By screening a comprehensive collection of E. coli mutations, we found that 45 *E. coli* mutations activate a *C. elegans* mitochondrial stress response gene *hsp-6*. But surprisingly, four of these *E. coli* mutations that affect enterobactin siderophore utilization or import into the *E. coli* cytoplasm induce a dramatic inviability of multiple *C. elegans* mitochondrial mutants. These *E. coli* siderophore mutations also caused a mild disruption of mitochondrial morphology in wild type and the induction of *C. elegans hsp-6* (Fig. 3A-B). But disruption of *E. coli* enterobactin biosynthesis did not cause this synthetic lethal interaction with a *C. elegans* mitochondrial mutants and could even suppress the synthetic lethality of *E. coli* mutants defective for enterobactin siderophore utilization or import into the cytoplasm, suggesting that the production of enterobactin is required for the toxic interaction between the iron loaded *E. coli* siderophore and *C. elegans* mitochondrial mutants.

We found that four of the mitochondrial mutations we tested were synthetic lethal with the enterobactin iron retrieval mutants but that the *gas-1* and *mev-1* mutants which also produce more ROS were not synthetic lethal. It may be important to the genetic interaction with the *E. coli* siderophore iron release mutations whether the ROS is generated in complex I or complex III, and how accessible it might be to the iron-loaded enterobactin. The detailed cryoEM structure of the entire animal mitochondrial ETC points to a highly intricate system of eight proteins bearing iron sulfur cluster redox centers that pass electrons with quantum mechanical precision, and efficiently couple to movements of transmembrane segments of other coupled Complex I proteins (there are 14 core proteins in complex I) to pump protons against a proton gradient (31). The many FeS clusters mediate the movement of electrons along the many protein subunits of complex I are the likely source of ROS in the complex I mutants. Interestingly, from the set of ETC mutants that are synthetic lethal with the four *E. coli* siderophore iron retrieval mutants, *nduf-7* encodes an orthologue of human NDUFS7 (NADH:ubiquinone oxidoreductase core subunit S7) which has a 4 Fe 4 S cluster The complex III ETC mutation *isp-1* also shows a strong synergistic interaction with the *E. coli* siderophore mutations. ISP-1 is an orthologue of human UQCRFS1 (ubiquinol-cytochrome c reductase, Rieske iron-sulfur polypeptide 1) and has a 2 iron, 2 sulfur clusters and ubiquinol-cytochrome-c reductase activity. Again, this is an iron sulfur center mutant protein that interacts with ubiquinone that shows such a strong synergy.

The *C. elegans spg-7* point mutation causes a mitochondrial defect and is strongly synthetically lethal with the *E. coli* siderophore Fe retrieval mutations. *spg-7* does not encode an ETC component, but encodes a protease that is the ortholog of the *E. coli* ftsH protease. This protease has a small number of client proteins, which includes the iron sulfur assembly factor iscS and the iron sulfur protein lpxC and fdoH. *C. elegans* mitochondrial proteins from the nuclear genome that are probable clients of SPG-7 are likely to be orthologs of the *E. coli* clients of ftsH: The *C. elegans* iron sulfur assembly factor *iscu-1* is a prime candidate for causing the iron sulfur protein defect in the ETC (32).

Similarly, the *clk-1*/mammalian coenzymeQ7 mitochondrial 5-demethoxyubiquinone hydroxylase that is synthetic lethal with the *E. coli* siderophore iron retrieval mutants fails to synthesize ubiquinone with its normally 10 isoprene unit hydrophobic tail in *C. elegans*. This null allele is not lethal to *C. elegans* because *E. coli* ubiquinone is imported into *C. elegans* cells. But the *E. coli* ubiquinione while competent for viability has a shorter bacterial isoprene tail and causes slow growth of *C. elegans*, probably due to inefficient transfer of electrons from complex I to *E. coli* ubiquinone or from *E. coli* ubiquinone to complex III. The most likely explanation for the synergistic lethality of the *C. elegans clk-1* mutation with the *E. coli* siderophore mutations is that the efficiency of *E. coli* ubiquinone interaction with complex I or complex III is compromised so that ROS to synergize with the iron loaded siderophore is generated (33).

Of the ETC subunits that do not interact with the *E. coli* siderphore mutations includes *gas-1* and *mev-1. gas-1* is orthologous to human NDUFS2 (NADH:ubiquinone oxidoreductase core subunit S2) does not have an iron sulfur cluster. *mev-1* is an ortholog of human SDHC (succinate dehydrogenase complex subunit C) and is a heme protein. *mev-1* is component of complex II, which does not participate in the NADH to oxygen redox/proton pumping cascade and would not be expected to generate ROS under normal redox conditions.

We propose that ROS production at particular sites in the ETC is the key to the genetic interaction with *E. coli* siderphore mutations. Such localized production of ROS may not be detectable when measuring ROS levels in extracts; but the localized ROS within complex I or III could explain the phenotype. In fact, the free radical scavenger suppression, where such small scavenger molecules may weave into Fe S clusters is evidence that free radicals are involved.

It is important to note that all of the mitochondrial mutations used in this study are non-null point mutations. The null phenotypes for all these genes are lethal (based on gene inactivations by RNAi). Thus, the complexities of which mitochondrial mutations synergize and which do not can be interpreted based on the functions of each protein, but these interpretations are diminished by the fact that single point mutations were screened (ie, these are not null phenotypes). Still, the correlation with Fe S proteins and *E. coli* siderophore genetic interactions is striking. And the ability of antioxidants to suppress the arrest induced by *E. coli* mutants defective in enterobactin siderophore utilization or import with *C. elegans* mitochondrial mutants suggests a ROS-based mechanism for the toxicity. Because endogenous ROS production is minimal in wildtype *C. elegans*, whereas mitochondrial mutants have increased ROS production, we propose that the triggering ROS in the *C. elegans* mitochondrial mutant animals when combined with ferric-chelated enterobactin induces a much larger ROS burst via the Fenton reaction to cause toxicity to *C. elegans*. However, in wildtype *C. elegans*, since there is minimal ROS production, there is not enough ROS to initiate the Fenton reaction; thus *E. coli* enterobactin siderophore utilization or import mutants do not induce larval arrest in wild type *C. elegans*. Nevertheless, the ferric chelated enterobactin that enters the wildtype *C. elegans* does activate the mitochondrial stress response. Another possibility is that since the mitochondria in the specific mitochondrial mutants are mildly dysfunctional, adding the toxic ferric chelated enterobactin exacerbates the mitochondrial damage. Since the mitochondria of the wildtype *C. elegans* are healthier, the *E. coli* enterobactin utilization or import mutants do not cause larval arrest in wildtype. In summary, our findings highlight the importance of gut microbial metabolites in regulating animal health through conserved mitochondrial bioenergetics pathways.

The interaction between bacterial siderophore iron retrieval mutations and *C. elegans* electron transport chain components that we have discovered may be grounded in the complex ecology of bacterial siderophores. Siderophores allow bacteria to retrieve the vanishingly small concentrations of soluble Fe(III) in the environment, including in cells (34). While many bacterial species synthesize siderophores, unrelated bacteria hijack the iron-bound siderphores from those siderophore synthesizing species to usurp their iron, a limiting essential micronutrient. But some bacterial species with siderophore biosynthetic pathways have evolved siderophore biosynthetic modifications to append antibiotics to the siderphore to inhibit the growth of these siderphore-iron thieves (34). In fact, enterobactin has been engineered to carry antibiotics as a sort of Trojan Horse for antibiotic delivery (35). Such an enterobactin-biotin fusion chemical was used to search for *C. elegans* proteins that interact with enterobactin, detecting the ATP synthase alpha subunit ATP-1 that is also homologous to atpA of bacteria (36). Although ATP synthase is not an iron-rich protein, it does cross the same mitochondrial membrane and uses the pH gradient generated by the electron transport chain to power the ATP synthase rotary motor. The iron-rich Complex I of the ETC passes electrons along the redox gradient from NADH, eventually via Complexes III and IV to oxygen, coupling that energy to the movements of iron-sulfur and heme proteins to pump protons. A 21 amino acid peptide of the ATP-1 ATP synthase alpha subunit that is sufficient to bind enterobactin (36), is highly conserved in the bacterial kingdom as well as in eukaryotes (Fig. S7). The proton flux from the ATP synthase rotary motor protein could react with the Fe bound to enterobactin to activate its ROS amplification from the complex I and complex III Fe S protein mutations that we have documented here. Thus, the *E. coli* enterobactin-Fe can be retrieved by diverse bacteria that may not have the proper enzymes (such as Fes) to remove the Fe so strongly bound to multiple catecholamines of enterobactin; this potentially toxic role of enterobactin-Fe may be a highly evolved role to target the universal components of bacterial and eukaryotic mitochondrial electron transport chains.

## Acknowledgments

We thank members of the G.R. laboratory for helpful discussions. The work was supported by NIH Grant AG043184 (to G.R.). **Some strains were provided by the CGC, which is funded by NIH Office of Research Infrastructure Programs (P40 OD010440)**.

## Author contributions

J.A.G., E.J, and G.R. conceived the project. J.A.G. performed all experiments with technical help from E.J. J.A.G. analyzed all the data, made the figures, and wrote the paper. G.R. supervised the experiments and wrote the paper with the help from the other authors.

## Declaration of Interests

The authors declare no competing interests.

