## Supplementary Information for "ROS-based lethality of *C. elegans* mitochondrial electron transport mutants grown on *E. coli* siderophore iron release mutants"

Supplementary Information Text

**Supplemental Materials and methods**

***C.elegans* strains used in this study**

The following strains were used in the study:

| Wildtype | N2 |
| --- | --- |
| SJ4100 | *zcIs13[hsp-6::GFP]* |
| SJ4058 | *zcIs9[hsp-60::GFP]* |
| SJ4005 | *zcIs4 [hsp-4::GFP] V* |
| AU133 | *agIs17 [myo-2p::mCherry + irg-1p::GFP] IV* |
| YL162 | *atfs-1(tm4525); zcIs9[hsp-60::GFP]* |
| QC115 | *atfs-1(et15) V* |
| DA2249 | spg-7 (ad2249) I |
| MQ130 | clk-1(qm30) III |
| CW152 | gas-1(fc21) X |
| MQ887 | isp-1(qm150) IV |
| TK22 | mev-1(kn1) III |
| CU5991 | fzo-1(tm1133) |
| CU6372 | drp-1(tm1108) |
| QC134 | nduf-7(et19) |
| VC1789 | nduf-2.2(ok2397) |
| MQ1333 | nuo-6(qm200) |
| NL1832 | *ucr-2.3(pk732)* |

**Genomewide keio screen for developmental delay or *hsp-6::gfp* induction phenotype**

Keio mutants which represent ~3800 genes were grown in LB media containing kanamycin at 37^o^C overnight. 100μl of the overnight culture were dropped onto NGM media containing 24-well plates and grown overnight at 37^o^C. Synchronized L1-stage transgenic *hsp-6::gfp* animals prepared by bleach-prep were dropped onto these 24-well Keio mutant in NGM media plates. The wells were screened visually at day 3 under a fluorescent microscope for the induction of *hsp-6::gfp*. All the wells were screened visually at day 5 under the DIC microscope for wells that contained animals growing significantly slower compared to control wells grown on wildtype BW25113 *E. coli*.

The primary hits were rescreened again at least three times.

**Removal of Kanamycin cassette in Δ*fes::Kan* Keio mutant and construction of double mutants**

Because bacterial insertion drug resistant cassette mutations in one gene in an operon are expected to be polar on downstream genes, the drug resistant cassette alleles are best for identifying relevant operons rather than individual genes. But because genes in operons often act in the same pathway, the identification of the operon is informative (Table S1). Mutations that disrupt the *E. coli* cytochrome oxidase *cyoABCDE* operon were previously shown to induce *C. elegans* *hsp-6:GFP* (1) ; these were among the strongest hits of the 24 mutants that caused developmental delay (Table 1). None of the other *E. coli* genes identified in the screen of the Keio mutant collection act in the *E. coli* electron transport chain, consistent with our previous finding that inactivation of only specific steps in bacterial respiration causes defects in *C. elegans* growth or mitochondrial function (1). Keio mutants have a Kanamycin cassette flanked by FLP recombinase sites, which allows for the excision of the cassette in the place of gene of interest (2). Kanamycin cassette incision in *Δfes::Kan* was achieved by introducing the plasmid pCP20 by transformation, which contains an Flp recombinase. Transformants were selected for ampicillin resistance at 30^o^C and the raised at 42^o^C to remove the pCP20 plasmid. Colonies were verified by inability to grow on LB plates with Kanamycin as well as by sequencing for the recombination site junction.

Double mutants, *Δfes entA::kan*, *Δfes entB::kan*, *Δfes cirA::kan*, *Δfes fiu::kan*, or *Δfes fepA::kan* were generated as follows: P1 lysates of Keio collection mutants, *entA::kan*, *entB::kan*, *cirA::kan*, *fiu::kan*, or *fepA::kan* strains were generated by growing overnight cultures of each strain in LB media containing 2mM MgSO_4_,10mM CaCl_2_, 0.2% glucose at 37^o^C until the OD_600_ is around 0.5. 100 μl of Phage P1 obtained form CGSC was added to 1 ml of each culture. After 2 hours incubation at 37^o^C until complete lysis, 0.5 ml of chloroform was added the culture. The lysates were centrifuged at 13K RPM for 1 min and the lysates were stored in 4^o^C until use. 100 μl of P1 lysate from each of the Keio mutant was added to overnight cultures of *Δfes* mutants grown in LB was suspended in 1ml LB media containing 5mM MgSO_4_, 50mM CaCl_2_ and incubated at 37^o^C for 1 hour. 0.5 ml of 10mM Sodium citrate was added to each tube and incubated for 30 min. The tubes were centrifuged at maximal speed for 1 min and plated on kanamycin containing LB Media agar plates. The colonies were verified for the presence of *Δfes* and Keio mutant insertions using sequencing.

***In vitro* ATP measurement**

ATP measurements were conducted as per the protocol with the following modifications (3). 1000 animals in 100 μl of M9 were frozen in liquid nitrogen and stored at -80°C until analysis. The samples were placed immediately in boiling water bath for 15 min. After adding twice amount of distilled water, centrifugation was done at 14,000 rpm for 5 min and supernatant is transferred into a second tube. Equal volume of luciferase reagent (Promega^TM^) was added and luminescence was measured using a plate reader (SpectraMax Microplate reader). Protein content was measured using BCA method. ATP concentrations were normalized to protein content of samples. All the values were normalized to wildtype grown on *E. coli* BW25113.

***In vivo* ATP measurement**

The transgenic reporter strain PE254 *(*[*feIs4*](http://www.wormbase.org/db/get?name=WBTransgene00000494;class=Transgene)*[Psur-5::luc+::gfp; rol-6 (su1006)] V)* (4) was used as a surrogate for determining the intracellular ATP content. ~1000 synchronized PE254 animals grown in *E. coli* BW25113 or *Δfes* mutant until the L4-larval stage were collected from NGM media plates and washed at least 4 times in M9 buffer to remove bacteria. The animals were transferred to white flat-bottom, 96-well plate wells containing 100 μM D-luciferin in M9 buffer and the luminescence was measured within 5 min in SpectraMax microplate reader. GFP fluorescence was quantified in SpectraMax microplate reader using 485nm excitation and a 520nm emission filter. Background measurements were subtracted from readings. Luminescence values were normalized to GFP fluorescence of samples. All the values were normalized to wildtype grown on *E. coli* BW25113.

**Measuring endogenous hydrogen peroxide levels using HyPer strain**

To measure endogenous hydrogen peroxide levels, transgenic *jrIs1*[P*rpl-17*::HyPer] animals expressing a HyPer as described (5). Synchronized L1-stage transgenic *jrIs1*[P*rpl-17*::HyPer] animals were grown in *E. coli* BW25113 or *Δfes* mutant and about 1000 L4-stage animals were harvested in 96 microtiter well plates. For H_2_O_2_ treatment experiment, H_2_O_2_ was added to the animals in the 96-well itself and measurements were taken within 15 minutes. An excitation wavelength of either 490 nm or 405 nm was used to measure oxidized or reduced HyPer probe fluorescence respectively with an emission filter of 535 nm.  The absorbance at 620 nm was used to normalize for animal numbers. The statistical significance of differences between conditions was determined by using unpaired *t*-test. GraphPad Prism 8.0 was used for these calculations.

**Staining of mitochondria**

Animals were stained with 10μM of 10-nonyl acridine orange bromide (NAO) in M9 solution for 1 hour and destained by washing in M9 atleast 5 times.

**Antioxidant treatments**

Stock solutions of trans-resveratrol, L-ascorbic acid, cis-resveratrol, oxyresveratrol, cis-trimethoxyresveratrol, trolox, or N-acetyl cysteine were diluted in M9 to a working concentration of 10μg/ml and dropped onto NGM media plates seeded with overnight cultures of *Δfes* mutant *E. coli*. Synchronized L1-stage *spg-7(ad2249)* animals were added to the plates and scored after three days for the suppression of arrest phenotype. The negative control NGM plates contained the solvent and M9.

**Microscopy and imaging**

Nematodes were mounted onto agar pads and images were taken using a Zeiss AXIO Imager Z1 microscope fitted with a Zeiss AxioCam HRm camera and Axiovision 4.6 (Zeiss) software. Student’s t test was used determine statistical significance.

Low-magnification bright-field images were acquired using a Zeiss AxioZoom V16 Microscopre, equipped with a Hammamatsu Orca flash 4.0 digital camera camera, and using Axiovision software. Some Immunofluorescence images were acquired with an IX-70 microscope (Olympus, Waltham, MA) fitted with a cooled CCD camera (CH350; Roper Scientific) driven by the Delta Vision system (Applied Precision, Pittsburgh, PA). Images were deconvolved using the SoftWoRx 3.3.6 software (Applied Precision).

**Quantification and Statistical analysis**

Statistical analysis was performed with GraphPad Prism 7 software. Student’s t-test was used for analysis.

**Fig. S1**

**Fig. S1: Screen methodology**

Individual Keio mutant *E. coli* strains were grown in individual wells of 24 well NGM agar plates, which supports bacterial growth on the surface, and the bacteria that grew on the agar surface were inoculated the next day with ~30 synchronized L1 larval stage *hsp-6::gfp* animals. Each individual well was then screened at two different time-points: 48 hours at 20^o^C for the Keio mutant strains that induced *hsp-6::gfp* and at 96 hours at 20^o^C for *E. coli* mutant strains that dramatically slowed *C. elegans* development.

Fig. S2

**Fig. S2: *E. coli* mutants induce *hsp-6::gfp***

While *hsp-6::gfp* is not induced in animals grown on wildtype *E. coli* BW25113, animals grown on mutant Keio *E. coli* strains showed *hsp-6::gfp* activation.

Fig. S3

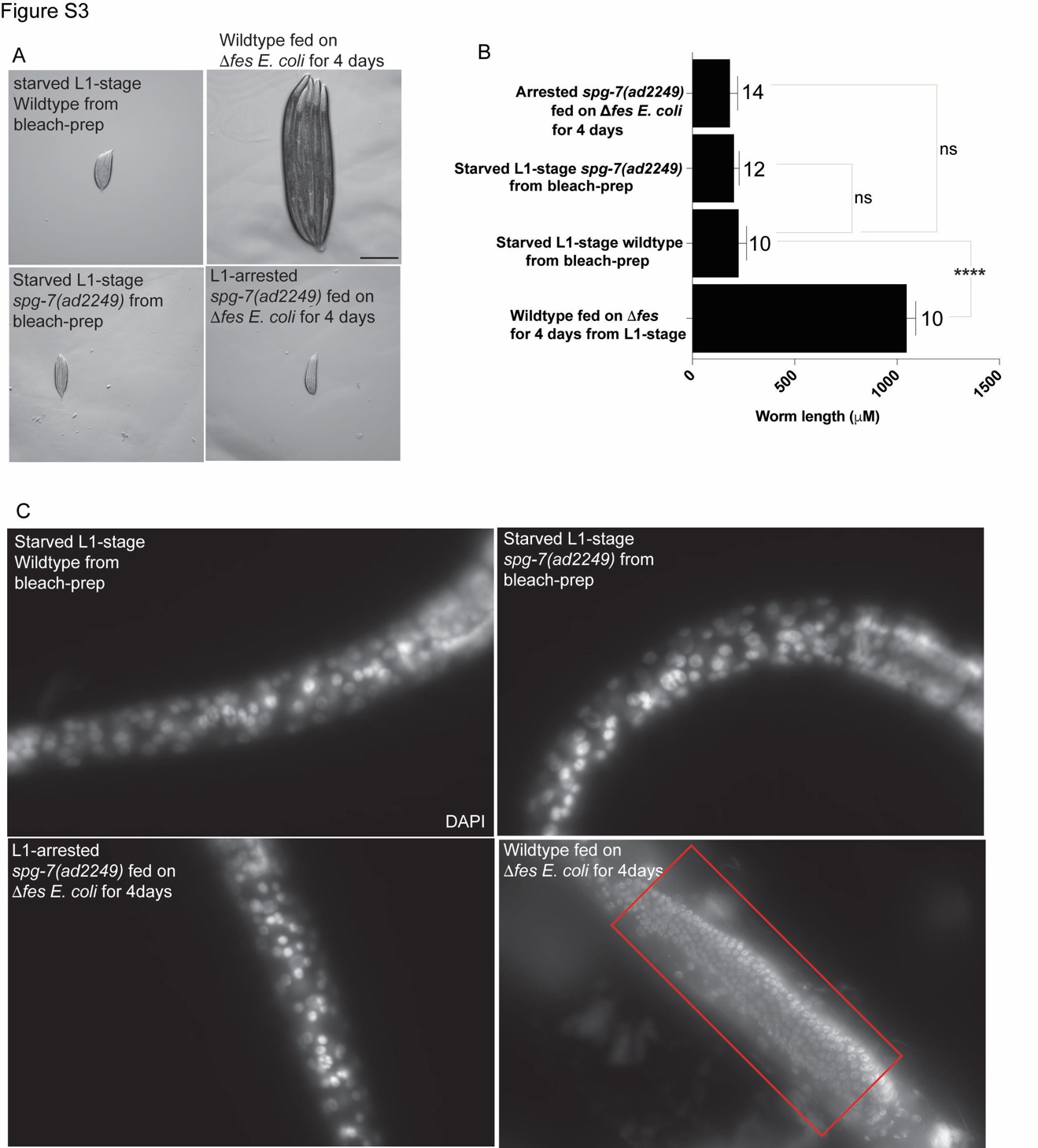

**Fig. S3: Determining the arrest stage of *spg-7(ad2249)* animals grown on Δ*fes-722* mutant**

A) The size and appearance of L1-larval stage wildtype or *spg-7(ad2249)* animals from egg-prep looks very similar to that of the arrested *spg-7(ad2249)* animals grown on *fes-722* mutant *E. coli* for 4 days. One-day old wildtype animals used for comparison was grown on *fes-722* mutant *E. coli* from L1-stage for 4 days. Scale bar, 100μM.
B) Quantification of the animal length in L1-larval stage wildtype or *spg-7(ad2249)* animals from egg-prep and arrested *spg-7(ad2249)* animals grown on *fes-722* mutant *E. coli* for 4 days. One-day old wildtype animals used for comparison were grown on *fes-722* mutant *E. coli* from L1-stage for 4 days. Statistical significance was determined using one-way ANOVA. **** P<0.0001. ns denotes not significantly different. The number of animals used is shown above each bar in the graph. The error bars indicate standard deviation. Data represent one out of two independent experiments.
C) Whole-mount DAPI staining of animals from L1-larval stage wildtype or *spg-7(ad2249)* animals from egg-prep and arrested *spg-7(ad2249)* animals grown on *fes-722* mutant *E. coli*. One-day old wildtype animals were used for comparison. The germline in the L1-larval stage wildtype or *spg-7(ad2249)* animals from egg-prep and arrested *spg-7(ad2249)* animals grown on *fes-722* mutant *E. coli* were not apparently visible at this stage because at this stage there are only 2 germline progenitor cells. If the arrested worms were in their L2 or L3 stages they would have apparently visible germ cells. The germline from adult wildtype animal (red box) is shown for comparison.

Fig. S4

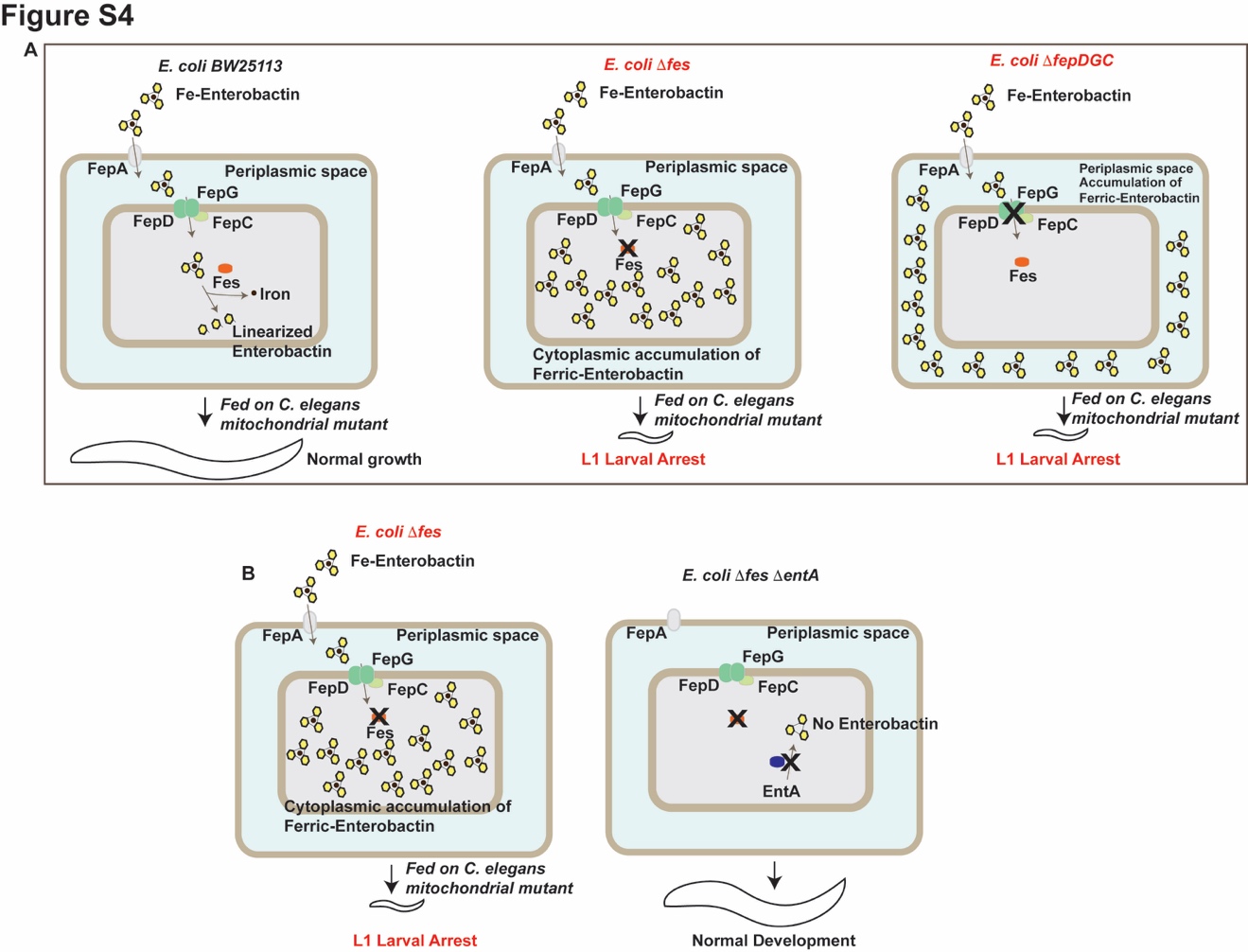

**Fig. S4: Diagrammatic representation of bacterial enterobactin content**

A) Diagrammatic representation of *Δfes::kan*, *ΔfepD::kan*, *ΔfepG::kan*, *ΔfepC::kan* mutant accumulating ferric enterobactin
B) Diagrammatic representation of *Δfes* mutant accumulating ferric-enterobactin and lack of enterobactin in *Δfes ΔentA::kan* double mutants.

Fig. S5

**Fig. S5: Interaction between mitochondrial mutants and Δ*fes-722*** **mutant**

*E. coli* mutants cause development arrest phenotype in *nduf-7(et19)*, *isp-1(qm150)*, and *clk-1(qm30)* mutants. No development arrest phenotype was observed in *gas(fc21)* and *ucr-2.3(pk732)* mutants grown on Δ*fes-722*.

Fig. S6

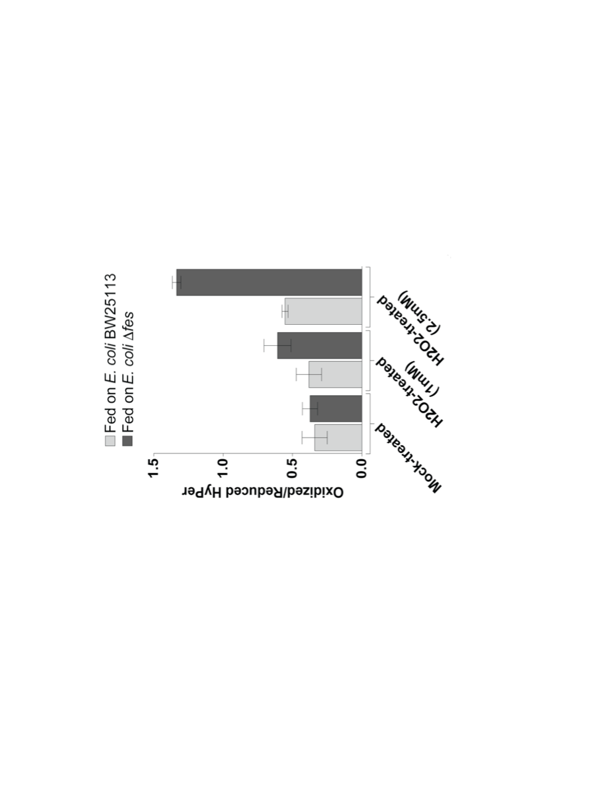

**Figure S6: Measuring hydrogen peroxide levels**

Endogenous hydrogen peroxide levels were not significantly different between animals grown on *E. coli* BW25113 or *Δfes-722 E. coli*; however, addition of exogenous peroxide results in significantly higher level of endogenous peroxide levels in animals grown on *Δfes-722 E. coli* compared to animals grown on *E. coli* BW25113.

Fig. S7

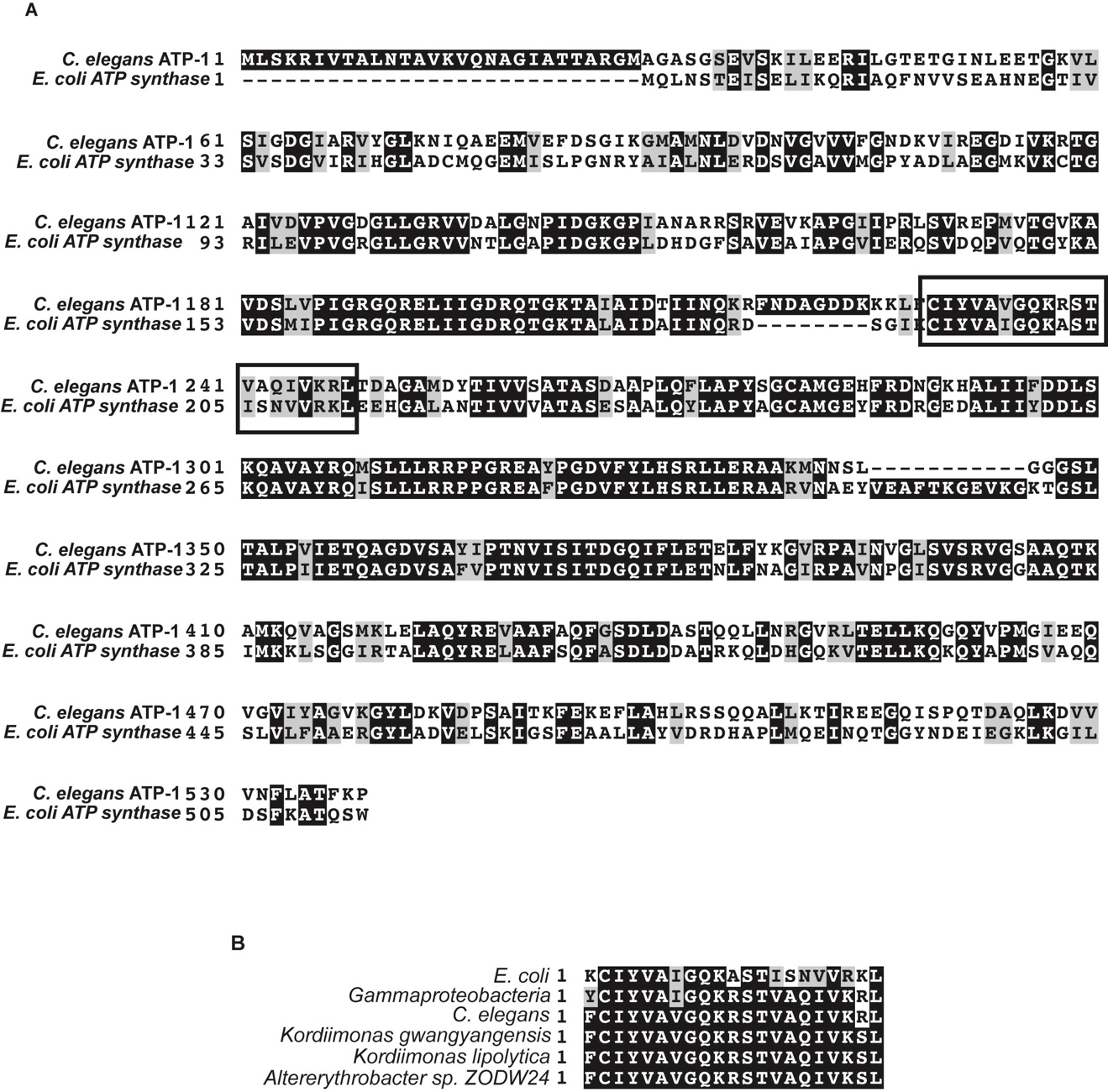

**Figure S7: ATP Synthase protein sequence analysis**

A) C. elegans ATP-1 is highly homologous to *E. coli* ATP synthase. The black boxes shows the 21 amino acid peptide sequence that was shown to bind enterobactin
B) The 21 amino acid peptide sequence shown to bind enterobactin is highly conserved

**Table S1: *spg-7(ad2249)* animals grown on Δ*fes-722* mutant *E. coli* killed with gentamicin or tetracycline arrest as L1-larvae.**

|  | ***spg-7(ad2249)* % Larval arrest (N)** |
| --- | --- |
| Fed on BW25113 | 0±0 (161) |
| Fed on Δ*fes-722*::*Kan* | 100±0 (188) |
| Fed on BW25113, Ampicillin-treated | 0.5±0.8 (176) |
| Fed on Δ*fes-722*::*Kan*, Ampicillin-treated | 0±0 (146) |
| Fed on BW25113, Tetracycline-treated | 0±0 (144) |
| Fed on Δ*fes-722*::*Kan*, Tetracycline-treated | 98.6±1.1 (155) |
| Fed on BW25113, Gentamicin-treated | 0.7±1.2 (144) |
| Fed on Δ*fes-722*::*Kan*, Gentamicin-treated | 98.4±1.7 (195) |
| Fed on BW25113, Kanamycin-treated | 0±0 (134) |
| Fed on Δ*fes-722::Kan*, Kanamycin-treated | 97.7±2.2 (175) |
| Fed on Δ*fes-722::Kan* + BW25113 (75:25) | 0±0 (119) |
| Fed on Δ*fes-722::Kan* + BW25113 (75:25), Kanamycin-treated | 98±1.9 (151) |
| Fed on Δ*fes-722::Kan* + BW25113 (50:50) | 0.65±1.1 (146) |
| Fed on Δ*fes-722::Kan* + BW25113 (50:50), Kanamycin-treated | 97.8±2.4 (142) |
| Fed on Δ*fes-722::Kan* + BW25113 (25:75) | 0±0 (138) |
| Fed on Δ*fes-722::Kan* + BW25113 (25:75), Kanamycin-treated | 98.7±2.1 (166) |
| Fed on Δ*fes-722::Kan* + BW25113 (100:1) | 0±0 (141) |
| Fed on Δ*fes-722::Kan* + BW25113 (100:1), Kanamycin-treated | 99±1.6 (166) |
| Fed on Δ*fes-722::Kan* + BW25113 (1000:1) | 0±0 (152) |
| Fed on Δ*fes-722::Kan* + BW25113 (1000:1), Kanamycin-treated | 98.5±2.4 (140) |
| Fed on Δ*fes-722::Kan* + BW25113 (10,000:1) | 0±0 (124) |
| Fed on Δ*fes-722::Kan* + BW25113 (10,000:1), Kanamycin-treated | 99.4±0.9 (163) |

Data from three independent trials were collected; the average of percentage of L1 animals and the standard deviation are shown. The total number of animals counted in all the three independent trials is shown in the parentheses.

**Table S2: Measurement of CFU/ml of wildtype *E. coli* BW25113 and *E. coli* *Δfes-722::Kan* in NGM plates**

|  | BW25113 (CFU/ml) | fes::Kan (CFU/ml) | Ratio |
| --- | --- | --- | --- |
| Day 0 | 1x10^4^ | 1x10^8^ | 10,000 |
| Day 1 | 2.6x10^7^ | 5.6x10^8^ | ~200 |
| Day 2 | 6x10^8^ | 4.6x10^8^ | ~1.3 |

**Table S3: Table showing the *spg-7(ad2249)*, *nduf-7(et19)*, *isp-1(qm150)*, *clk-1(qm30)* tested on N-Acetyl Cysteine antioxidant for suppression of arrest phenotype**

|  | **% Larval arrest (N)** | |
| --- | --- | --- |
| **Animal Strain** | **Fed on *E. coli* *Δfes-722;* Solvent control** | **Fed on *E. coli* *Δfes-722;* NAC** |
| Wildtype | 0±0 (142) | 0±0 (161) |
| *spg-7(ad2249)* | **100±0 (147)** | 0.69±1.2 (151) |
| *nduf-7(et19)* | **100±0 (142)** | 1±1.8 (129) |
| *isp-1(qm150)* | **100±0 (121)** | 0±0 (108) |
| *clk-1(qm30)* | **100±0 (155)** | 0.7±1.3 (144) |

Data from three independent trials were collected; the average of percentage of L1 animals and the standard deviation are shown. The total number of animals counted in all the three independent trials is shown in the parentheses.

**Table** **S4: Trans-resveratrol treatment acts through antagonizing animal ROS pathways**

| ***spg-7(ad2249)* animals fed on Δ*fes-722*** | **% Larval arrest (N)** |
| --- | --- |
| solvent control | **100±0 (105)** |
| Trans-Resveratrol, 10μg/ml | 0±0 (123) |
| Trans-Resveratrol, 10μg/ml, washed | **100±0 (143)** |

*spg-7(ad2249)* animals grown on Δ*fes-722* mutant grown in trans-resveratrol arrest do not arrest while animals grown on Δ*fes-722* mutant *E. coli* pre-grown in the presence of trans-resveratrol and washed to remove trans-resveratrol arrest as L1-larvae.

**Supplementary Dataset 1**

**Table showing list of the hits from the keio screen for mutants that induces *hsp-6::gfp*. All these mutant were found to induce *hsp-60::gfp*. None of these mutants activated *hsp-4::gfp* or *pgp-5::gfp*.**
